# When does accounting for gene-environment interactions improve complex trait prediction? A case study with *Drosophila* lifespan

**DOI:** 10.1101/2025.07.12.664531

**Authors:** Fabio Morgante, Francesco Tiezzi

## Abstract

Gene-environment interactions (G × E) have been shown to explain a non-negligible proportion of variance for a plethora of complex traits in different species, including livestock, plants, and humans. While several studies have shown that including G × E can improve prediction accuracy in agricultural species, no increase in accuracy has been observed in human studies. In this work, we sought to investigate the reasons for the contradictory results between agricultural species and humans. Model organisms are useful for studying G × E, since environments can be controlled and genotypes can be replicated across environments. Thus, we used data from a previous study in *Drosophila melanogaster*, where the authors measured lifespan in different environments and found evidence of G × E. We devised three different cross-validation (CV) scenarios that mimic the relationships between reference and test populations observed in agriculture and human studies, and fitted a few statistical models with and without including G × E. The results showed that G × E explained 8% of lifespan variance. Despite that, models accounting for G × E improved prediction accuracy only in CV scenarios where the same genotypes are observed in both the reference and test populations. While these scenarios are common in agriculture, where individuals of the same family or variety appear in both populations, they are not encountered in human studies, where individuals are unrelated. Thus, our work clarifies in which prediction scenarios we can expect improvements by incorporating G × E into statistical models, and provides an explanation for previous results found in studies involving human populations.

## Introduction

Estimating the effect of gene-environment interactions (G × E) on complex traits and accounting for it in phenotype prediction is one of the most difficult challenges faced by plant and animal breeders [1]. Recently, this topic has also become of interest in human genetics [2–7].

In plant breeding, the estimation of such interaction largely relies on multi-environment trials [8]. Here, some varieties of a given species are planted in multiple locations, managed under different agronomic techniques and exposed to different pedo-climatological conditions. The estimates of gene-environment interaction will be non-null when the performance of a given genotype varies across the different locations [8]. This implies that the genotype is sensitive to environmental conditions in its expression. Plant breeders leverage this interaction for developing cultivars that thrive in the intended conditions or, perhaps, across conditions [9]. Similarly, animal breeders use datasets where related individuals (*e*.*g*., paternal half-sibs) are raised in different environments, so to estimate how the same or similar genotypes can acclimate to different conditions [10, 11]. This is done mostly to breed for animals that can be tolerant to stress (*e*.*g*., heat stress), but also to develop “the right genotype for that environment”, like ruminants that can adapt well to grazing conditions [12].

Many statistical approaches for plant and animal breeders have been developed [13]. These approaches can be summarized as: 1) multiple trait models (MTM) that consider the trait of interest as a series of correlated traits, each defined by the ‘environment’ under which is manifested; 2) random-regression models (RRM) that model the change in the performance across conditions (*i*.*e*., reaction norm models), with conditions described by one or a few environmental covariates; and 3) Reproducing Kernel Hilbert Space regression (RKHS) models that can handle a large number of covariates in interaction with genotypes. All these models have been described in the literature, and the relationships between them can be demonstrated, at least in theory [1, 13–15]. It should be noted that MTM is the least parsimonious in terms of parameters to estimate when the number of environments or environmental descriptors becomes large. On the other hand, RKHS is the most parsimonious, as it was developed specifically to handle a large number of environmental covariates [14].

Models that incorporate G × E have shown good promise in plant and animal breeding, although the magnitude of the benefit is strongly dependent on the structure of the training population. In plants, Jarquin *et al*. showed an increase in modeling the performance of wheat lines grown across dozens of locations [16], while Acosta *et al*. successfully modeled general and specific combining ability in interaction with the environment for predicting the performance of maize hybrid lines [17]. In addition, Cuevas *et al*. showed that G × E implemented in RKHS models produced advantages with non-null correlations between environments [18]. In livestock, both Bohlouli *et al*. and Tiezzi *et al*. found a clear advantage in modeling G × E using climate data in dairy cattle, using random RRM or RKHS models, respectively [19, 20].

In human genetics, researchers cannot design optimal experiments for G × E analysis and need to rely on large observational data such as biobanks [21]. These data have a large amount of missing records, rely on noisy self-assessed information about the environment, lack replication of genotypes (*i*.*e*., individuals) across environments, and genotypes are non-randomized with respect to environments [5]. This results in large difficulties in estimating the magnitude of G × E. Despite that, non-negligible contribution of G × E to the phenotypic variance of several complex traits has been shown [2, 6, 22]. However, accounting for G × E has generally not resulted in an increase in out-of-sample prediction accuracy [3, 5, 7, 23]. These results disagree with those obtained in agricultural breeding.

To study in which scenarios G × E is expected to improve accuracy and explain the contradictory results between humans and agricultural species, model organisms can provide valuable insights. Experiments in model organisms can be highly controlled, resulting in precise environment definitions and replication of genotypes across such environments, which is ideal for G × E analyses. In this study, we used data from an experiment that measured lifespan for flies from the *Drosophila melanogaster* GeneticReference Panel (DGRP) raised in different environments [24, 25]. Using these data, we sought to investigate the conditions in which accounting for G × E in different statistical models improves out-of-sample prediction accuracy.

## Materials and methods

### Data processing

We used phenotypic data from [25]. Lifespan (in days) was measured for 186 inbred lines of the DGRP for the two sexes in three different temperatures (18°C, 25°C, 28°C). After removing lines that had missing values in at least a sex/temperature combination, we were left with *n* = 176 lines, each measured in *c* = 2 context variables (*r* = 6 sex/temperature combinations), for a total of *q* = 1, 056 records. The genotype data were filtered to remove genetic variants with minor allele frequency (MAF) smaller than 0.05 and missing genotype rate greater than 0.2. These filters retained *p* = 1, 899, 439 genetic variants.

### Statistical models

We analyzed the data using RKHS regression models described in [14]:

- G-BLUP. *y*_*ijk*_ = *µ* + *a*_*i*_ + *ϵ*_*ijk*_.
- E-BLUP. *y*_*ijk*_ = *µ* + *e*_*ij*_ + *ϵ*_*ijk*_.
- GE-BLUP. *y*_*ijk*_ = *µ* + *a*_*i*_ + *e*_*ij*_ + *ϵ*_*ijk*_.
- G×E-BLUP. *y*_*ijk*_ = *µ* + *a*_*i*_ + *e*_*ij*_ + *ae*_*ij*_ + *ϵ*_*ijk*_.

where

*y*_*ijk*_ is the phenotype of line *i* in environment *j*,

*µ* is the intercept value,

*a*_*i*_ is the random additive genetic value of line 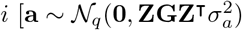, **Z** is a *q* × *n* incidence matrix, **G** is a *n* × *n* genomic relationship matrix (GRM) computed as in [26]],

*e*_*ij*_ is the random environmental value of line *i* in environment 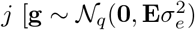, **E** is a *q* × *q* matrix of similarity based on environmental variables, computed as **E** ∝ **XX**^⊺^, **X** is a *q* × *c* matrix of environmental measurements],

*ae*_*ij*_ is the random gene-environment value of line *i* in environment *j* 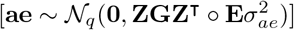,

*ϵ*_*ijk*_ is the residual value for line *i* in environment 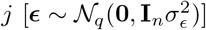.

These models were fitted in a Restricted Maximum Likelihood (REML) framework, implemented in the `sommer R` package [27].

We also fit a MTM [28]:

- mvG-BLUP. **Y** = **M** + **A** + **R**

where

**Y** is an *n* × *r* matrix of phenotypic observations in the *r* = 6 environments,

**M** is an *r*-vector of intercept values,

**A** is an *n* × *r* matrix of additive genetic values [**A** ∼ MN _*n×r*_(**0, G, Σ**_*A*_), **Σ**_*A*_ is an *r* × *r* genetic covariance matrix],

**R** is an *n* × *r* matrix of residual values [**R** ∼ MN _*n×r*_(**0, I**_*n*_, **Σ**_*R*_), **Σ**_*R*_ is an *r* × *r* diagonal residual covariance matrix].

This model was fitted in a Bayesian framework, implemented in the `BGLR R` package [29]. We ran the sampler for a total of 300,000 iterations, discarding the initial 200,000 iterations as burn-in, followed by thinning every 50 iterations.

We also fit a RRM [30]:

- RRM. 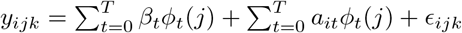

where

*y*_*ijk*_ is the phenotype for line *i* in environment *j*,

*ϕ*_*t*_(*j*) is the Legendre polynomial of order *t* for environment *j*,

*β*_*t*_ is the fixed effect of the *t*^*th*^-order Legendre polynomial,

*a*_*it*_ is the random additive genetic value of the *t*^*th*^-order Legendre polynomial [**a** ∼ N_*q*_(**0, G** ⊗ **Σ**_*a*_), **Σ**_*a*_ is a *T* × *T* genetic covariance matrix],

*ϵ*_*ijk*_ is the residual value for line *i* in environment *j* [***ϵ*** ∼ N_*q*_(**0, I**_*n*_ ⊗ **Σ**_*ϵ*_), **Σ**_*ϵ*_ is an *r* × *r* diagonal residual covariance matrix whose *j, j* element is 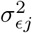]. We chose T=1 and used the mean phenotype across lines within each environment as the environmental value to compute *ϕ*_*t*_(*j*). This model was fitted in a Restricted Maximum Likelihood (REML) framework, implemented in the sommer R package [27].

### Validation Schemes

We implemented three cross-validation (CV) schemes, whereby part of the data (*i*.*e*., the training set) was used to train the models and the remaining part (*i*.*e*., the test set) was used to evaluate prediction accuracy.

1. *Random Lines*. We assigned 17% of the lines (for all the combinations of sex and temperature) to the test set, randomly (Fig 2A). This procedure was repeated 6 times. The peculiarity of this scheme is that the lines in the test set are not represented in the training set. Thus, the training-test transfer of information happens mostly at the environmental level.
2. *Random Observations*. We assigned 17% of the observations (*i*.*e*., combinations of line, sex, and temperature) to the test set, randomly (Fig 2B). This procedure was repeated 6 times. The peculiarity of this scenario is that all the lines, sexes, and temperatures are represented in the training set. Thus, the training-test transfer of information happens at both the genetic and environmental level.
3. *New Environment*. We assigned all the observations in a specific combination of sex and temperature to the test set (∼ 17% of the data) (Fig 2C). This procedure was repeated 6 times. The peculiarity of this scenario is that a sex/temperature combination is never seen in the training set. Thus, while technically the training-test transfer of information happens at both the genetic and environmental level, for the latter, it is hampered by not observing the actual environment we are trying to predict in.

Prediction accuracy was computed as *R*^2^ from the regression of the true phenotypes on the predicted phenotypes, averaged over the 6 replicates.

## Results and Discussion

We first partitioned the phenotypic variance into sources of variation attributed to genetics, environment, and gene-environment interactions (Fig 1A).

**Fig 1.**
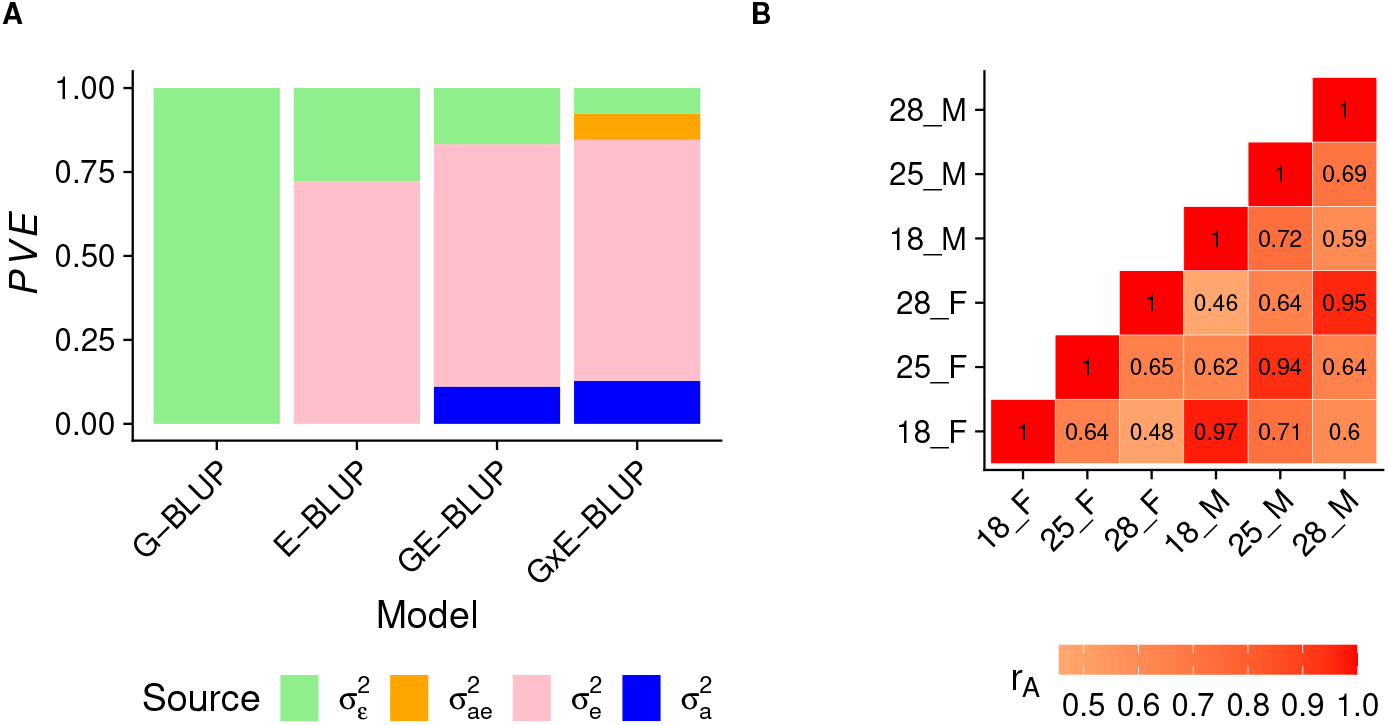
Variance partition and cross-environment genetic correlations of lifespan. A) Variance partition into genetic 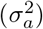, environmental 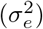, gene-environment interaction 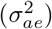, and residual 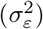 components using RKHS models. The y-axis shows proportion of variance explained (PVE). B) Cross-environment genetic correlations (*r*_*A*_) estimated using MTM.

The results (Fig 1A) show that the environment explained the largest amount of lifespan variance (∼ 72%) across all models that include that component. The genetic variance was the second largest contributor in GE-BLUP and G × E-BLUP, explaining ∼ 12% of the variance. However, genotypes explained no variance in G-BLUP. We attributed this result to the fact that when the environment explains most of the variance and its effect is not modeled, the model struggles to find the (minor) genetic signal in the large amount of unexplained variance. But once the effect of the environment is included in GE-BLUP and G × E-BLUP, it is easier for genetic effects to explain some of the remaining variance (*i*.*e*., not accounted for by the environment). Gene-environment interactions explained ∼ 8% of the phenotypic variance. Importantly, when including gene-environment interactions in the model, they explained variance that would be otherwise included in the residual, as implied by the similar proportion of variance explained (PVE) by genetic effects and environmental effects in GE-BLUP and G × E-BLUP. We also sought to confirm the presence of gene-environment interactions in a complementary analysis, where we estimated cross-environment genetic correlations (*r*_*A*_), treating lifespan in each of the six environments (*i*.*e*., sex/temperature combinations) as a different trait in mvG-BLUP [28]. The results (Fig 1B) show that the genetic correlations are different from unity for every pair of environments, especially across temperatures. The observation that genetic effects are different across environments agrees with the presence of gene-environment interactions [31].

We then assessed the accuracy of the different models at predicting yet-to-be-observed phenotypes using three CV schemes, illustrated in Fig 2.

**Fig 2.**
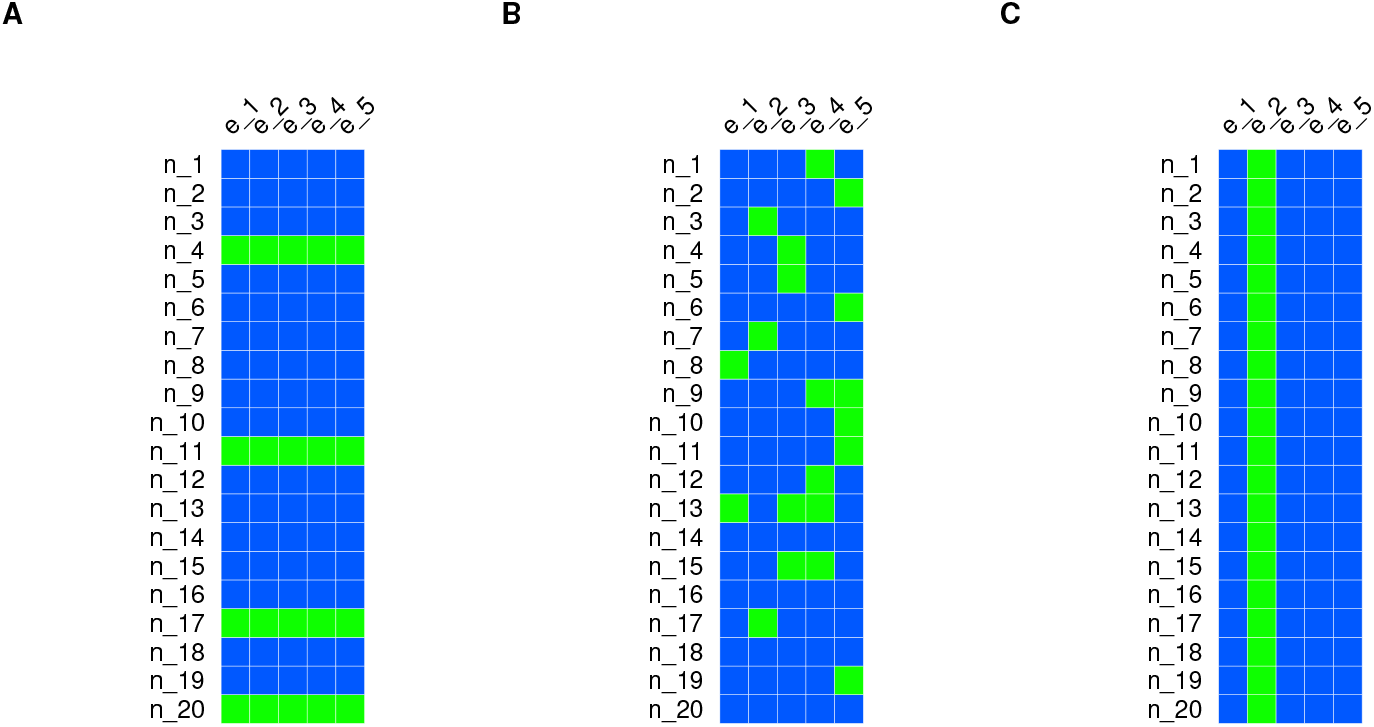
Cross-validation schemes used to evaluate the prediction accuracy of the different models. A) *Random Lines*. B) *Random Observations*. C) *New Environment*. The blue squares represent observations in the training set and the green squares represent observations in the test set.

The prediction results are shown in Fig 3.

**Fig 3.**
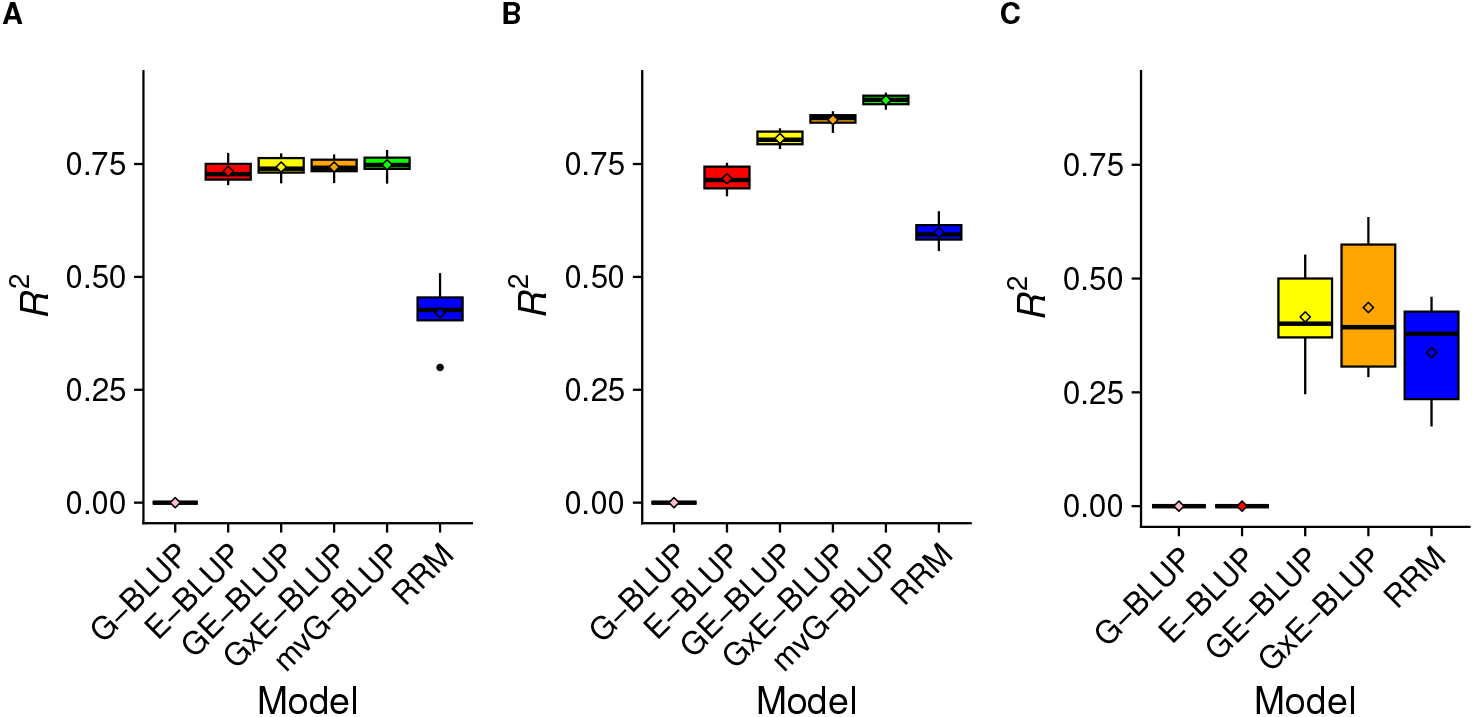
Prediction accuracy in the different cross-validation schemes. A) *Random Lines*. B) *Random Observations*. C) *New Environment*.

In the *Random Lines* scenario (Fig 3A), G-BLUP performed poorly. This result is expected, since the lines assigned to the test were not present in the training set (and DGRP lines are largely unrelated [24]) and G-BLUP was shown to explain little variance in the whole dataset. On the other hand, E-BLUP had high prediction accuracy (*R*^2^ ≈0.72). Here, sex and temperature were shown to explain ∼ %72 of the variance and all the sex/temperature combinations were observed in the training set, making the estimation of their effects an easy task. The prediction accuracy provided by GE-BLUP and G × E-BLUP was similar to (although slightly higher than) that of E-BLUP, showing that including genetic effects did not help improve predictions much in this scenario. mvG-BLUP also has similar accuracy to E-BLUP, GE-BLUP, and G × E-BLUP. This result stems from the fact that mvG-BLUP fits lifespan in each environment as a different trait and thus estimates environment-specific intercepts, genetic and residual variances. The difference in phenotype due to the different environmental conditions is accounted for by the intercepts in mvG-BLUP. The percentage of variance explained by this model ranged ∼ 72 − 80% across environments, showing that genetic effects could explain a large proportion of within-environment phenotypic variance. mvG-BLUP also leveraged the medium-high cross-environment genetic correlations to achieve good prediction accuracy. On the other hand, RRM had a much lower accuracy compared to the other environment-aware models. These results are in partial disagreement with other studies that showed RRM to be competitive with the other models [32]. An explanation for this observation is that our data might be suboptimal for the application of RRM, since the environments appear to be better defined as discrete rather than stratified on a continuous scale.

In the *Random Observations* scenario (Fig 3B), prediction accuracies were generally higher than in the *Random Lines* scenario. This is expected since all the lines and environments were present in the training set – only specific line/environment *combinations* were not observed in the training set. Thus, *Random Observations* is an easier scenario than *Random Lines* for prediction. G-BLUP performed poorly and LUP did well (*R*^2^ ≈ 0.72) again. However, GE-BLUP improved accuracy substantially over E-BLUP, providing *R*^2^ ≈ 0.81 in this scenario. This shows that once the large proportion of variance explained by the environment is accounted for, genotypes can explain additional variance and improve prediction accuracy in this scenario where all the genotypes (and environments) are observed in the training set. Accounting for gene-environment interactions in G × E-BLUP increased accuracy further (*R*^2^ ≈ 0.85), showing that this full model can achieve high accuracy at predicting specific genotype/environment combinations when all genotypes and all environments are observed in the training set. The best performing model was mvG-BLUP, which achieved *R*^2^ ≈ 0.89. This is a remarkable result since *R*^2^ from this model closely approached the PVE of the full model (*i*.*e*., G × E-BLUP) in the whole dataset (∼0.92). Thus, a model that explicitly accounts for the specificity of genetic effects across environments, while also leveraging their similarities, seems to be the best choice.

However, this model has some limitations: (1) it requires discrete, well defined environments, with a low to moderate number of classes; (2) it conflates the contributions of genetics and gene-environment interactions to the phenotypic value into a single term; (3) it does not allow for the prediction in unobserved environments, unless one has estimates of the genetic and residual (co)variances between observed and unobserved environments, and the phenotypic mean in the unobserved environments. Again, RRM had a much lower accuracy compared to the other environment-aware models.

The *New Environment* scenario (Fig 3C) is the most difficult among our CV schemes, which resulted in much lower prediction accuracies than in the other scenarios. In fact, the environment in which we are trying to predict is not observed in the training set, and the environmental conditions explain the majority of the phenotypic variance. This is reflected in E-BLUP performing poorly in this scenario. However, GE-BLUP could achieve moderate accuracy (*R*^2^ ≈0.42), presumably due to a better disentanglement of genetic and environmental effects when estimating parameters in the training set. G × E-BLUP provided slightly improved accuracy (*R*^2^ ≈0.44) compared to GE-BLUP, confirming the utility of accounting for G×E when predicting in unobserved environments. In this scenario, mvG-BLUP could not be fitted because of reason (3) discussed in the previous paragraph. While RRM performed worse than GE-BLUP and G × E-BLUP on average, its accuracy (*R*^2^ ≈ 0.34) was much closer to the accuracies of those models in this scenario. In the *New Environment* scenario, there were substantial differences in prediction accuracy depending on which environment the prediction was made (Table 1).

**Table 1.** Prediction accuracy (*R*^2^) in each environment (Sex_Temperature) in the *New Environment* cross-validation scenario.

| Environment | G-BLUP | E-BLUP | GE-BLUP | G×E-BLUP | RRM |
| --- | --- | --- | --- | --- | --- |
| F_18°C | 0.00 | 0.00 | 0.55 | 0.63 | 0.37 |
| M_18°C | 0.00 | 0.00 | 0.52 | 0.63 | 0.44 |
| F_25°C | 0.00 | 0.00 | 0.37 | 0.37 | 0.17 |
| M_25°C | 0.00 | 0.00 | 0.43 | 0.42 | 0.19 |
| F_28°C | 0.00 | 0.00 | 0.25 | 0.29 | 0.46 |
| M_28°C | 0.00 | 0.00 | 0.37 | 0.28 | 0.39 |

Overall, at least one of the models accounting for gene-environment interactions (*i*.*e*., G × E-BLUP and RRM) performed as well as or better than GE-BLUP in every environment. As expected, given the estimates of the cross-environment genetic correlations, differences in prediction accuracy were larger among temperatures than between sexes (within temperature). At 18°C, G×E-BLUP greatly outperformed GE-BLUP. At 25°C, G×E-BLUP and GE-BLUP yielded similar accuracy. RRM performed substantially worse than the other methods at both 18°C and 25°C. On the other hand, at 28°C, RRM achieved higher accuracy than GE-BLUP and G× E-BLUP.

Our results showed that accounting for gene-environment interactions can be helpful for improving predictions. However, the presence and magnitude of the improvement depends on the prediction scenario. In general, predicting the performance of certain genotypes in certain environments largely depends on the phenotyping strategy for building reliable training populations [16]. If the goal is to predict phenotypes for unrelated individuals in known environments (our *Random Lines* scenario), accounting for G × E will not improve prediction accuracy. A potential explanation for this observation could be the lack of information sharing at the genetic level between the training set and the test set, coupled with the additional parameters that need to be estimated in models accounting for G× E.

At the other end of the spectrum, if the goal is to predict phenotypes for a set of known individuals in a new environment (our *New Environment* scenario), accounting for G × E will most likely improve prediction accuracy. A plausible explanation for this observation is that G × E-aware models can predict the adaptability of each genotype to several conditions. This is particularly relevant for agricultural breeding, where breeders need to know how available breeds/cultivars would fare in a new environment [33, 34]. However, the choice of the modeling method is key here, with some G × E-aware methods outperforming others depending on which environment we aim to predict in.

Finally, if the goal is to predict unobserved phenotypes for a set of known individuals in known environments (our *Random Observations* scenario), that is the scenario where we can expect the largest increase in accuracy when accounting for G × E. The *Random Observations* scenario is particularly relevant to precision medicine, where we are interested in predicting medically relevant phenotypes (*e*.*g*., blood pressure) after a change in the environment (*e*.*g*., switching from a high fat diet to a low fat diet). In this scenario, a multivariate model treating phenotypes in different environments as different traits seemed to provide the highest accuracy. However, this scenario cannot be evaluated with human data, as individuals are present in only one level of the environmental variable (*e*.*g*., a person either smokes or does not, but not both). This peculiarity might also explain why accounting for G × E – despite explaining non-negligible variance – has generally not resulted in improved prediction accuracy for human traits [3, 5, 7]. In fact, assigning some individuals to the test set and trying to predict their phenotype in a known environment (as done in human studies) is equivalent to our *Random Lines*: in this scenario, we have shown that including G × E does not improve predictions.

Our study has some important limitations. First, the DGRP has a small sample size, limiting the accuracy with which genetic effects can be estimated. Second, all the models used in this study do not perform variable selection, implying that all genetic variants, all environmental variables, and all the interactions between them have an effect on lifespan. Using methods that perform variable selection has the potential to increase prediction accuracy further. Third, our study focused on only one trait, as this was the only one available for a well designed and controlled experiment, which allowed us to avoid confounding effects. Thus, our results will need to be confirmed on additional traits from equally well designed, but larger experiments.

## Conclusion

In this study, we have shown in which scenarios we can expect gene-environment interactions to improve phenotypic prediction accuracy, providing an explanation for previous results of human studies.

## Data availability

The genotype data are available at http://dgrp2.gnets.ncsu.edu/. The phenotype data are available from [25]. Code used for the analyses is available at https://github.com/morgantelab/dgrp_lifespan_gxe.

